# Cooperative Contributions of Structural and Functional Connectivity to Successful Memory and Aging

**DOI:** 10.1101/144154

**Authors:** Simon W Davis, Amanda Szymanski, Homa Boms, Thomas Fink, Roberto Cabeza

**Affiliations:** Center for Cognitive Neuroscience, Duke University, Durham, NC, USA; Department of Neurology, Duke University School of Medicine, Durham, NC, USA.

**Keywords:** aging, connectivity, DWI, associative memory, item memory

## Abstract

Understanding the precise relation between functional connectivity and structural (white-matter) connectivity and how these relationships account for cognitive changes in older adults are major challenges for neuroscience. We investigate these issues using a new approach in which structural equation modeling (SEM) is employed to integrate functional and structural connectivity data analyzed with a common framework based on regions connected by canonical tract groups (CTGs). CTGs (e.g., uncinate fasciculus, cingulum, etc.) serve as a common currency between functional and structural connectivity matrices, and ensures that the same amount of data contributing to brain-behavior relationships. We used this approach to investigate the neural mechanisms supporting memory for items and memory for associations, and how they are affected by healthy aging. Our results are threefold. Firstly, structural and functional CTGs made independent contributions to associative memory performance, suggesting that both forms of connectivity underlie age-related changes in associative memory. Secondly, distinct groups of CTGs supported associative versus item memory. Lastly, the relationship between functional and structural connectivity was best explained by the relationship between latent variables describing functional and structural CTGs based on a constrained set of tracts—but no one specific CTG group—suggesting that age effects in connectivity are constrained to specific pathways. These results provide further insights into the interplay between structural and functional connectivity patterns, and help to elucidate their relative contribution to age-related changes in associative memory performance.

## Introduction

One of the most consistent patterns in the literature on episodic memory and aging is that older adults tend to be more impaired in episodic memory for associations than in episodic memory for individual items. While this behavioral dissociation has been well known for a long time (Glisky et al., 1995; Naveh-Benjamin, 2000), cognitive neuroimaging provides a complementary method for investigating the underlying neural mechanisms (for review, see Old and Naveh-Benjamin, 2008). During the last three decades, cognitive neuroimaging has gradually moved from an emphasis on individual brain regions to a focus on the interactions among brain regions, or connectivity. Connectivity, which can be examined at the functional level using functional MRI (fMRI) and at the structural level using diffusion-weighted imaging (DWI). Given that functional connectivity depends on structural (white-matter) connectivity, a current challenge is how to investigate the relationship between these two forms of connectivity in relation to cognitive function. Here, we propose a new approach for linking structural and functional connectivity data and apply it to the results of an fMRI-DWI study investigating item and associative memory in younger and older adults.

There are two main challenges in linking structural and functional connectivity. First is the problem of *translation* between structural and functional information; structural matrices are considerably more sparse than functional networks (Wang et al., 2015), and while structural connectivity is static, functional connectivity is highly dependent on the active process concurrent with data collection (Honey et al., 2007). Second is the problem of the *granularity of mapping*; while a large array of techniques that have attempted to delineate structural-functional connectivity relationships at the level of whole-brain parcellations (Betzel et al., 2014; Zimmermann et al., 2016), between discrete pairs of regions (Andrews-Hanna et al., 2007; Dennis et al., 2008; Davis et al., 2012), or at the level of voxels (Horn et al., 2014) each technique tends to form a unique claim about how the structure-function relationship changes with age. Both of these problems preclude any lasting or satisfying conclusions about how these modalities relate to one another, and have issues unique to datasets that include older adults. The structural equation modeling (SEM) approach used here attempts to address these problems, and provides a rigorous statistical framework to examine the complex relationships between age, structural integrity of white matter, functional correlations between regional task-driven activity, and cognitive performance.

The premier method for assessing age-related changes in structural is DWI tractography. While the use of semi-automated pipelines for generating whole-brain connectomes based on DWI tractography has engendered an explosion of connectome-based research (for review, see Wang et al., 2015), the construction method varies widely across studies, and these pipelines are rarely informed by known anatomy, reducing their validity and replicability. This lack of analytical sensitivity and specificity may lead to a lack of consistency across construction methods (Thomas et al., 2014; Zhong et al., 2015), and contributes to a measure that is dominated by false positives (de Reus and van den Heuvel, 2013; Drakesmith et al., 2015). Despite this uncertainty, a number of anatomically defined, canonical white matter tracts demonstrate reliable relationships between white matter integrity and memory, including the fornix, uncinate fasciculus, cingulum, and the genu of the corpus callosum. Multiple indices of integrity of these PFC-based white matter tracts have been associated with age-related changes in scores on verbal associative memory (Davis et al., 2009; Kennedy and Raz, 2009; Bendlin et al., 2010; Voineskos et al., 2010), spatial- (Oberlin et al., 2016), object-based associative memory (Antonenko et al., 2016), as well as free recall of both visual and verbal information (Metzler-Baddeley et al., 2011; Metzler-Baddeley et al., 2012). While there is a wide array of approaches to addressing the overall influence of age in these studies (e.g., partial correlation, mediation, change scores, etc.), the consistency of these tract-specific relationships suggests a relative specificity to of specific connections to specific forms of memory. Furthermore, tract-specific dissociations such as these observations are not solely attributable to the strong neural declines associated with this region (Jack et al., 2002; Sullivan et al., 2006), as well as the role that more global declines in frontally-mediated executive functions play in observed episodic memory decline (Kievit et al., 2014; Kievit et al., 2016).

While DWI measures are able to characterize structural differences across aging, fMRI has been used to study age-related differences in regional co-activation (or connectivity) associated with memory function. In one of the first studies to examine age-related differences in functional connectivity associated with episodic (item) memory, Grady and colleagues (2003) found a ventral-to-dorsal shift in functional coupling between the hippocampus and activity in the rest of the brain during successful episodic encoding. While young adults demonstrated connectivity from a hippocampal seed to posterior sensory regions, older adults exhibited greater success-related functional coupling with the dorsolateral PFC and parietal cortex. Thus, older adults demonstrated a shift from posterior-to-anterior connectivity. Although this study focused on item memory, the finding has been replicated in associative memory encoding (Dennis et al., 2008), and several other studies have observed similar levels of plasticity in hippocampal connectivity across aging. Increased frontotemporal connectivity attributed to an age-related increase in top-down modulation has also been found in emotional and associative memory studies (Murty et al., 2009; St. Jacques et al., 2009; Addis et al., 2010). These studies suggest that frontotemporal tracts like the uncinate fasciculus, inferior fronto-occipital fasciculus and the fornix play a privileged role in mediating age-related cognitive decline. However, it is unclear how these region-to-region changes emerge in the context of a fully-connected system; significant increases or reductions in bivariate estimates may emerge as a function of subtler global changes in connectivity. More recently, the use of whole-brain, task-based connectomes estimated either from PPI- or beta-series correlation-based methods (Fornito et al., 2012; Geib et al., 2015), aim to address this ambiguity, but these methods have yet to be applied to older adults.

A major goal of connectome research is to discover whether, and how, the structural and functional networks of the brain are related — an active area with tremendous interest and wide ramifications in neuroscience. Increasingly, the widespread use of automated connection matrices has led to an explosion of computational solutions to this problem, typically by directly comparing connectivity matrices (Horn et al., 2014), predicting one modality from the other (Bowman et al., 2012; Abdelnour et al., 2014; Messé, 2015), joint analysis of structural and functional matrices (Honey et al., 2009; Tewarie et al., 2014), or through the comparison of graph-theoretical properties common to structural and functional networks (Betzel et al., 2014; Romero-Garcia et al., 2014). These more data-driven approaches have produced a number of meaningful observations (for an excellent review, see Zhu, 2014), principally that the relationship between functional connectivity and structural connectivity also appears to increase across the lifespan, and that this relationship is driven by an increase in the reliance on more long-distance interactions between brain regions (Betzel et al., 2014; Meunier et al., 2014). Nonetheless, these computational approaches have largely ignored canonical divisions in the structural anatomy of human white matter pathways. This is problematic because in the case of structural models, these models rarely incorporate known anatomy, leading to spurious connections and a high false-positive rate in structural connectomics generally (Maier-Hein et al., 2016). In the case of functional information, these computational solutions rarely take into account the relative sparsity of structural connection matrices compared to functional data. Thus, finding the adequate basis on which to make the comparison between these modalities is challenging.

The present analysis seeks to address these gaps in understanding by using task-based functional connectivity, and whole brain structural connectivity informed by classical white matter anatomy. These two data types are united in a common analytical framework in order to ask a specific question: *do functional and structural connectivity make independent contributions to memory in older adults?* The structural equation modelling (SEM) approach adopted here provides a framework to explore brain-behavior relationships. Particularly, we explore the possibility that function-structure relationships are best characterized by either specific linkages between data types for a given region, or instead reflect a general relationship shared by task-relevant regions (or tract groups). We test a model fitting structural and functional connectivity information summarized by Canonical Tract Groups (CTGs) in order to predict associative and item memory in younger and older adults. Only by better understanding the functional and neural bases of different types of memory can we develop future strategies to help with age-related memory loss.

## Methods

### Participants

Seventy-six adults—54 older adults (67.68 ± 6.9 y.o., age range 61-87 y.o.) and 22 younger adults (23.6 ± 3.5 y.a., age range 19-28 y.o.)—participated in the study. All individuals were screened for contraindications to MRI, and seven individuals were excluded because of scanner issues or poor structural or functional imaging quality, and two individuals did not complete the memory task, leaving N = 67 with complete data. Written consent was obtained for each participant and they received monetary compensation at the end of the study. All experimental procedures were approved by the Duke University Institutional Review Board.

### Associative Memory Task

#### Materials

Stimuli consisted of 440 normative English words with normative word frequencies in the lexicon of 5-15 per million, M = 8.8 (3.1), and had a mean length of M = 7.1 (2.3) letters. Unique study and test lists were randomly generated for each participant and words were assigned to the following conditions: item (180 words), associative (180), or item lures (80 words—presented only at retrieval as new words). At retrieval, there were four item tests lists, each consisting of 45 targets (old words) and 20 lures (non-studied words), and four associative test lists, each consisting of 45 studied words.

#### Encoding Task

Participants studied the words outside the scanner. Words were presented on a computer monitor in white font on a grey background for 3 s with a 1 s interval using Cogent (http://www/vislab.ucl.ac.uk/cogent_2000.php), a stimulus presentation software within MATLAB (www.mathworks.com). For half of the trials, participants made a “pleasant/unpleasant” judgement, and a “bigger/smaller than a shoebox” judgement for the other half. Half of the trials were repeated, with the same judgement; however, for the purpose of the present study, words that were repeated were collapsed into one condition for the subsequent fMRI analysis.

#### Retrieval Task

Approximately 15 min after the encoding phase, participants were tested for their memory of the studied words in the MRI scanner. Words were presented via a mirror in the scanner head coil and a rear projection system using a PC computer running Cogent. There were two retrieval conditions: Item memory and associative memory. In the item memory retrieval task, participants made new/old responses on a 4-point confidence scale. For the associative memory retrieval task, participants were asked to indicate what type of judgment they made earlier on a word on a 4-point scale: definitely pleasant/unpleasant, probably pleasant/unpleasant, probably bigger/smaller, definitely bigger/smaller. Four item and 4 associative runs were presented in consecutive blocks to minimize the effects of task switching. Retrieval stimuli were presented for 3 s, with a white crosshair presented for fixation during the inter-trial interval (ITI). Stimulus order and ITI jitter (range: 1-7 s) were determined by a genetic algorithm designed to maximize statistical efficiency and facilitate deconvolution of the hemodynamic response (Wager and Nichols 2003).

### MRI Acquisition & Analysis

The analytical pipeline is summarized in Figure 1. Participants were first scanned on a 3-T gradient-echo scanner (General Electric 3.0 Tesla Signa Excite HD short bore scanner, equipped with an 8-channel head coil). Coplanar functional images were acquired using an inverse spiral sequence (64 × 64 matrix, time repetition [TR] = 1700 ms, time echo [TE] = 31 ms, field of view [FOV] 240 mm, 37 slices, 3.8-mm slice thickness, 254 images). Using a spiral-in gradient-echo sequence: slice order = interleaved, matrix = 64^2^, FOV = 24 cm, TR=2000ms, TE=27ms, sections=34, thickness=3.8mm, interscan spacing = 0, flip angle = 60, SENSE reduction factor = 2). Following functional imaging, a high-resolution SPGR series (1-mm sections covering whole brain, interscan spacing=0, matrix = 256^2^, flip angle = 30, TR = 22 ms, TE = min full, FOV = 19.2 cm) was collected. Finally, DWI data were collected using a single-shot echo-planar imaging sequence (TR = 1700 ms, slices = 50, thickness = 2.0 mm, FOV = 256 × 256 mm^2^, matrix size 128 × 128, voxel size = 2 mm^3^, b value = 1000 s/mm^2^, diffusion-sensitizing directions = 25, total images = 960, total scan time = 5 min). The anatomical MRI was acquired using a 3D T1-weighted echo-planar sequence (matrix = 2562, TR = 12 ms, TE = 5 ms, FOV = 24 cm, slices = 68, slice thickness = 1.9 mm, sections = 248). Scanner noise was reduced with ear plugs, and head motion was minimized with foam pads. Total scan time, including breaks and structural scans, was approximately 1 h 40 min. Behavioral responses were recorded with a 4-key fiber-optic response box (Resonance Technology, Inc.), and when necessary, vision was corrected using MRI-compatible lenses that matched the distance prescription used by the participant.

**Figure 1.**
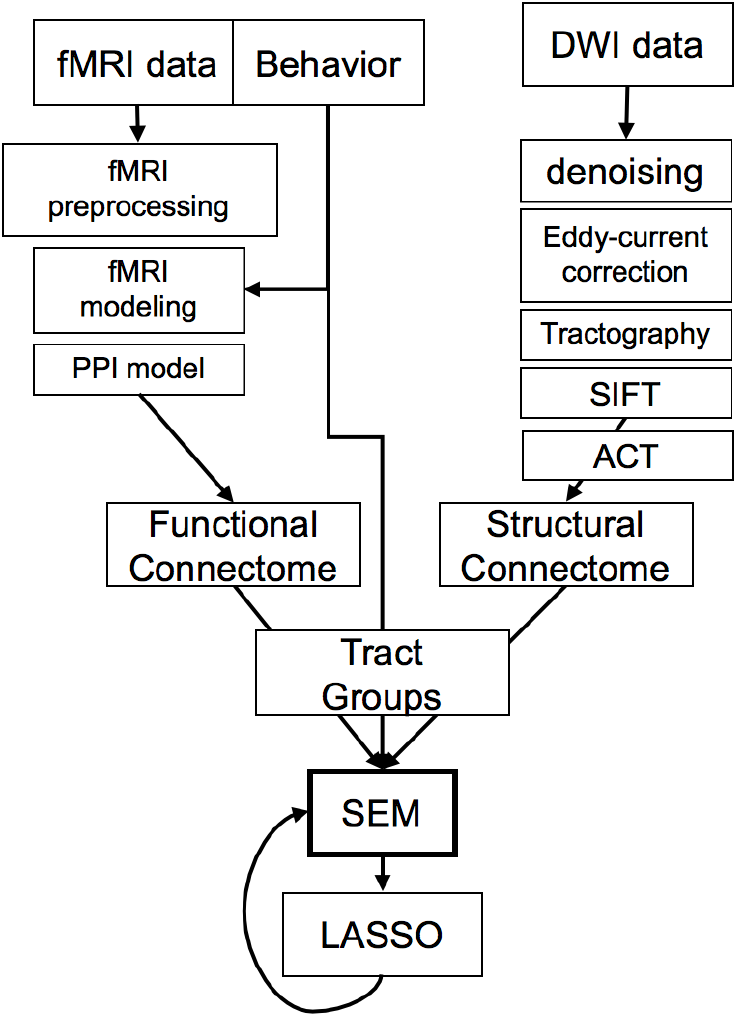
Analytical Pipeline.

#### Construction of connectivity matrices

Before either structural or functional matrices were constructed, we first sought to establish a consistent parcellation scheme across all subjects and all modalities (DWI, fMRI) that reflects an accurate summary of full connectome effects (Bellec et al., 2015). Subjects’ T1-weighted images were segmented using the SPM12 (<u>www.fil.ion.ucl.ac.uk/spm/software/spm12/</u>), yielding a grey matter (GM) and white matter (WM) mask in the T1 native space for each subject. The entire GM was then parcellated into 411 regions of interest (ROIs), each representing a network node by using a subparcellated version of the Harvard-Oxford Atlas, (Tzourio-Mazoyer et al., 2002), defined originally in MNI space. The T1-weighted image was then nonlinearly normalized to the ICBM152 template in MNI space using fMRIB’s Non-linear Image Registration Tool (FNIRT, FSL, <u>www.fmrib.ox.ac.uk/fsl/</u>). The inverse transformations were applied to the HOA atlas in the MNI space, resulting in native-T1-space GM parcellations for each subject. Then, T1-weigted images were coregistered to native diffusion space using the subjects’ unweighted diffusion image as a target; this transformation matrix was then applied to the GM parcellations above, using FSL’s FLIRT linear registration tool, resulting in a native-diffusion-space parcellation for each subject.

DWI data were analyzed utilizing FSL (https://fsl.fmrib.ox.ac.uk/fsl/fslwiki) and MRtrix (http://mrtrix.org) software packages. Data were denoised with MRtrix, corrected with eddy current correction from FSL, and brain extraction was performed with both FSL and MRtrix, whereas bias-field correction was completed with MRtrix. Constrained spherical deconvolution (CSD) was utilized in calculating the fiber orientation distribution (FOD). This FOD was used along with the brain mask to generate whole brain tractography, with seeding done at random within the mask (Tournier et al., 2004; Tournier et al., 2007). Relevant parameters regarding track generation are as follows: seed = at random within mask; step-size = 0.2 mm; 10,000,000 tracts. After tracts were generated, they were filtered using SIFT (spherical-deconvolution informed filtering of tractograms). This process utilizes an algorithm which determines whether a streamline should be removed or not based off of information obtained from the FOD, which improves the selectivity of structural connectomes by using a cost-function to eliminate false positive tracts (Yeh et al., 2016). Tracts were SIFTed until 1 million tracts remained. Prior to connectome generation, subject-specific MNI-space brains were created by an affine registration between the MNI T1 2mm brain template and b0s using FSL’s FLIRT. The MNI subject-specific brains then underwent another affine registration to the Harvard-Oxford 100 and 471 ROI templates. Once all registrations were completed, the connectomes were generated using the SIFTed tracts and the subject-specific templates for the 100 and 471 HOA images.

Functional connection matrices representing task-related connection strengths were estimated using a correlational psychophysical interaction (cPPI) analysis (Fornito et al., 2012). Briefly, the model relies on the calculation of a PPI regressor for each region, based on the product of that region’s timecourse and a task regressor of interest, in order to generate a term reflecting the psychophysical interaction between the seed region’s activity and the specified experimental manipulation. In the current study the task regressors based on the convolved task regressors from the univariate model described above were used as the psychological regressor, which coded subsequently remembered and subsequently forgotten word pairs with positive and negative weights, respectively, of equal value. This psychological regressor for successful memory retrieval was based on a linear contrast of Hits > Misses for both Associative and Item Memory blocks; new trials during the Item memory blocks were modeled, but not used in the connectivity analysis. This memory success-related regressor was multiplied with two network timecourses for region *i* and *j*. We then computed the partial correlation *ρ_PPI_i_,PPI_i_ · z_*, removing the variance *z* associated with the psychological regressor, the timecourses for regions *i* and *j*, and constituent noise regressors. We accounted for the potential effects of head motion and other confounds by assessing the 6 motion parameters and including these parameters in our partial correlation between regions.

#### Defining Tract Groups

We examine only region pairs that are connected by canonical fiber systems, which term here a Canonical Tract Group (CTG). This approach affords three main benefits, namely 1) integrating structural and functional connectivity information within a common anatomical framework, 2) constraining the overabundance of functional connections to known anatomy and 3) simplifying the number of pairwise comparisons in an informed manner. Tract group assignment is based on an overlap between matrix-based connections and canonical fiber systems from seven tracts defined by the Johns Hopkins University white matter tractography atlas (Hua et al., 2008): the uncinate fasciculus (UF), superior longitudinal fasciculus (SLF), inferior fronto-occipital fasciculus (IFOF), forceps minor (FMin), inferior longitudinal fasciculus (ILF), ventral cingulate gyrus (CINGhipp) and the dorsal cingulate gyrus (CING), as well as the fornix, based on a novel template (Brown et al., 2017). The corticospinal tract and forceps major were not included because they were not hypothesized to be involved in item or associative memory functioning.

Next, a region pair within a connectivity matrix in which the two regions share overlap with a given fiber tract are then considered part of the assembly for a CTG (Fig. 2). We refer to this matrix of n x n elements (where n is the number of regions) as the *CTG mask*. This assignment is then repeated for all 21 tract groups within the JHU Tract Atlas (FSL). Our analysis based on CTGs offers two clear advantages to data-driven comparisons between these data types: a) CTGs serve as a common currency between functional and structural connectivity matrices and b) this method addresses the fact that structural matrices are much more sparse than functional matrices. Thus, by using an identical set of region pairs from the adjacency matric in each tract group (e.g., the UF is shown in Fig. 2), we ensure that the same amount of data contributes to structural or functional connectivity information in the model.

**Figure 2.**
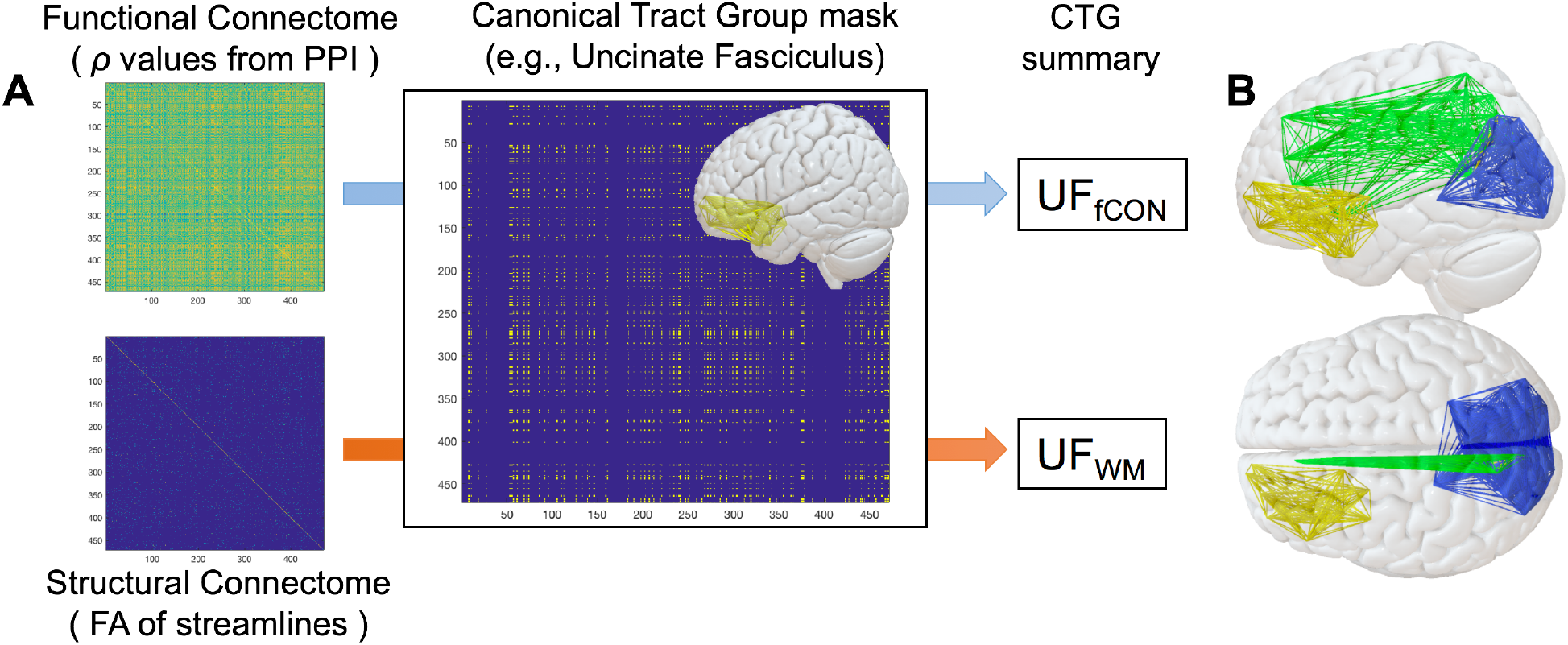
Construction of Canonical Tract Groups. **A**. After functional and structural connectome construction, a CTG mask for a particular tract is used to filter functional and structural connectivity information. All elements in the filtered matrix are then averaged create a functional and structural estimate for a given CTG (e.g., UF_fCON_ and UF_WM_) which is amenable to SEM modeling. Also obvious in these images is the relative sparsity of structural connection matrices comparted to functional matrices. **B**. Example CTGs for the uncinate fasciculus (yellow), cingulum (green), or forceps major (blue).

### Structural Equation Modeling

After Structural, Item, and Associative matrices filtered by CTGs and summed across all elements in the matrix, CTG values are averaged across hemisphere for bilateral tracts, and scaled before modeling. We fit confirmatory SEMs to the mean FA of the seven, bilaterally averaged, WM tract CTGs, which showed different sensitivities to age. These models were used to test the validity of three single-factor Latent Variables for structural connectivity (WM) based on DWI tractography, and functional connectivity (fCON) associated with successful retrieval, based on fMRI collected from either Item or Associative blocks. Both CFA and full SEMs were fit using the lavaan package (Rosseel, 2012) in R version 3.3.3 (R Development Core Team, 2016), and regularized SEM for complex models using regsem, (Jacobucci et al., 2016), which allows the use of LASSO-based regularization while keeping the SEM model intact, adding penalization directly into the estimation of the model. LASSO imposes a penalty on the regression parameters to ensure that the SEM model remains stable even when the number of predictors is large. Specifically, it uses the L1 norm to apply a least absolute shrinkage and selection operator (LASSO, Tibshirani, 1996) penalty, which enforces sparse solutions by shrinking many regression parameters to 0. We therefore applied LASSO regression to the 2 full model SEMs in order to penalize the models and reduce the number of contributing CTGs.

Prior to model fitting, variables were scaled to a standard normal distribution and log transformed where necessary to increase normality. All models were fit using Maximum Likelihood Estimation (ML) using robust standard errors and report overall model fit assessed with the chi-square estimates, Root Mean Squared Error of Approximation (RMSEA) and its confidence interval, the Comparative Fit index (CFI). We used the following guidelines for judging good fit (Bagozzi and Yi, 2012): RMSEA below 0.05 (acceptable: 0.05-0.08) and a CFI above 0.97 (acceptable: 0.95-0.97). Model comparison was estimated via the χ^2^ difference and the log likelihood ratio test. The significance of individual paths was tested with p-values less than 0.05, and the contribution of each predictor was assessed using the R^2^ value. Lastly, to test the influence of age (for both functional and structural CTGs) in the model, Age paths were set to zero, and model fit was reassessed using the same likelihood ratio test.

## Results

### Behavioral testing

Associative memory accuracy was 0.76 ± 0.016 and RTs were 2.09 s ± 0.49 for successful associative trials, and 2.37 s ± 0.58 for unsuccessful associative trials. Mean hit rates for item memory were 0.85 ± 0.015 and a mean false alarm rate of 0.22 ± 0.008. RTs during the item memory test were 1.59 ± 0.45 for item hits, and 2.36 for item misses. Effects of age were more prominent for associative memory than item memory, though the interaction between Age Group x Memory type was not significant (F_61,2_ = 2.18, p = 0.14). Effects of age for both associative memory (F61,1 = 12.45, p = 0.0008) and overall item memory hit rate (F_61,1_ = 3.68, p = 0.04) were significant.

### Canonical Tract Groups: Descriptive Statistics and Effects of Age

Following the method outlined above, we developed a semi-automated pipeline for assigning a given connection between ROIs within a standard atlas to a given Canonical Tract Group (CTG). While most CTGs showed significant age differences in the structural domain, success-related functional connectivity differences between younger and older adults were far subtler (Table 1). Functional CTGs showed consistently moderate relationships with corresponding structural CTGs (e.g., connectivity between regions functionally connected by the UF correlated with structural integrity of the UF), even after adjusting for the effects of age (all *r* > 0.21, p < 0.05).

**Figure 3.**
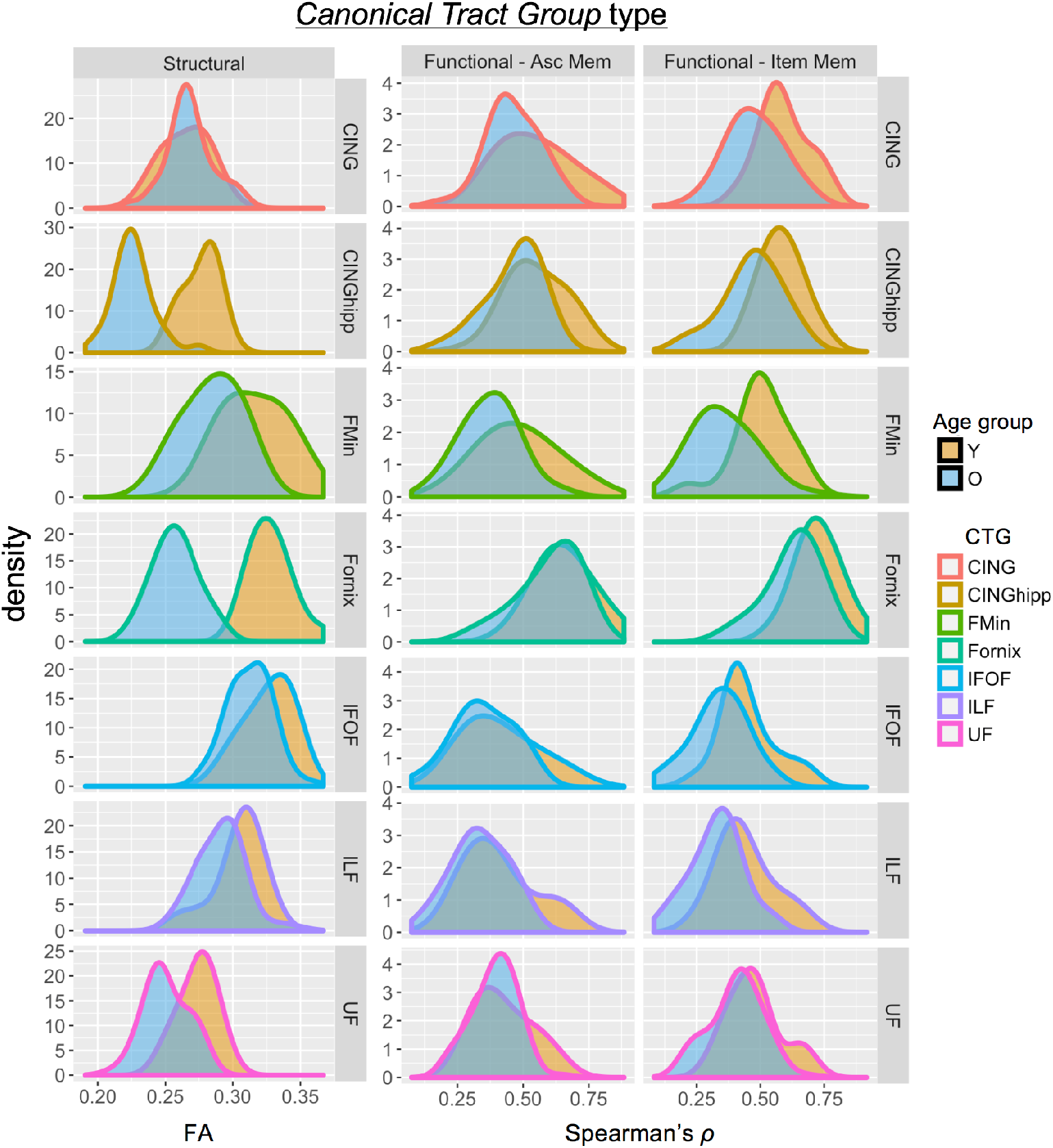
Age distributions of Canonical Tract Groups. Distribution of CTG values for structural connectivity (FA) and functional connectivity (Spearman’s rho, as calculated in PPI) for Associative and Item Memory in younger and older adults. CTG values are shown for seven major CTGs which influence the final SEMs below. Note: CING – cingulum; CINGhip – ventral leg of the cingulum; FMin – forceps minor (or genu); IFOF – inferior-fronto-occipital fasciculus; ILF – inferior longitudinal fasciculus; UF – uncinate fasciculus.

**Table 1.**
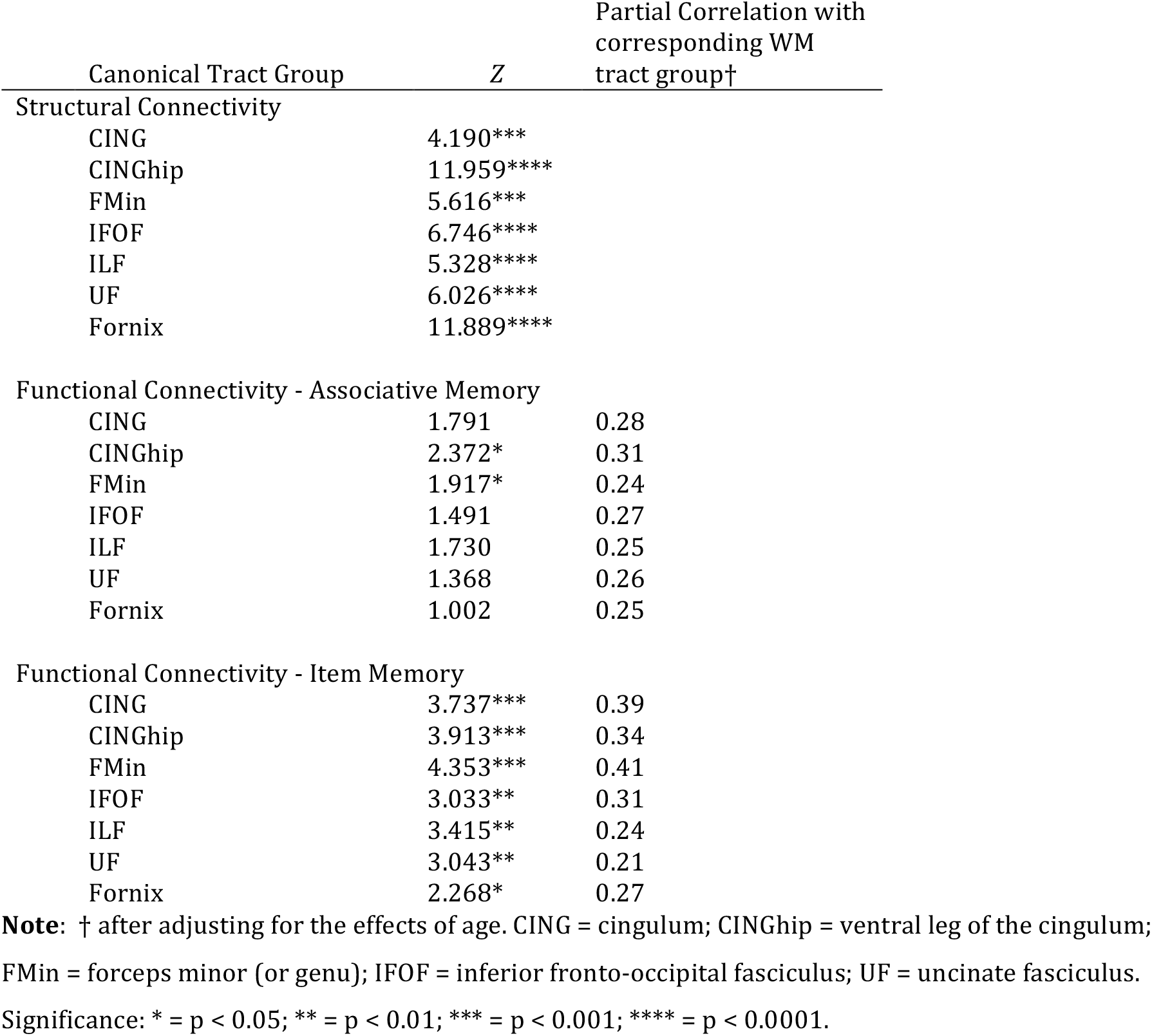
Effects of Age on Canonical Tract Groups.

| Canonical Tract Group | Z | Partial Correlation with corresponding WM tract group† |
| --- | --- | --- |
| Structural Connectivity |  |  |
| CING | 4.190*** |  |
| CINGhip | 11.959**** |  |
| FMin | 5.616*** |  |
| IFOF | 6.746**** |  |
| ILF | 5.328**** |  |
| UF | 6.026**** |  |
| Fornix | 11.889**** |  |
| Functional Connectivity - Associative Memory |  |  |
| CING | 1.791 | 0.28 |
| CINGhip | 2.372* | 0.31 |
| FMin | 1.917* | 0.24 |
| IFOF | 1.491 | 0.27 |
| ILF | 1.730 | 0.25 |
| UF | 1.368 | 0.26 |
| Fornix | 1.002 | 0.25 |
| Functional Connectivity - Item Memory |  |  |
| CING | 3.737*** | 0.39 |
| CINGhip | 3.913*** | 0.34 |
| FMin | 4.353*** | 0.41 |
| IFOF | 3.033** | 0.31 |
| ILF | 3.415** | 0.24 |
| UF | 3.043** | 0.21 |
| Fornix | 2.268* | 0.27 |

**Note:** † after adjusting for the effects of age. CING = cingulum; CINGhip = ventral leg of the cingulum;
FMin = forceps minor (or genu); IFOF = inferior fronto-occipital fasciculus; UF = uncinate fasciculus.
Significance: \* = $p < 0.05$ ; \*\* = $p < 0.01$ ; \*\*\* = $p < 0.001$ ; \*\*\*\* = $p < 0.0001$ .

### SEM results

#### CFA results

We used SEM to test a range of models of how connectivity supports associative memory in older adults. We first examined the reliability of the measures to be used in three confirmatory factor analysis (CFA) models; in these models, we hypothesize that two latent variables (functional connectivity or fCON and structural connectivity or WM) capture the covariance between 7 connectivity measures estimated from specific CTGs (described above), freely estimating every factor loading. Three CFA models were run: WM, fCON_item_, and fCON_associative_.

#### Full model results

Next, using CTGs we fit two full models relating brain connectivity variables to behavioral variables using a standard SEM. These models capture the hypothesis that individual differences in structural and functional connectivity measures make independent contributions to successful memory functioning. As a number of tracts and therefore CTGs should be linearly correlated, we can ask whether a more parsimonious model shows better fit.

We fit two models, one which focuses on Associative Memory, and one which focuses on Item Memory; while these models share some overlap in the regions demonstrating predictive power in the model, our use of LASSO allows for distinct tract groups to emerge as significant predictors in each model. The full model for Associative Memory is shown in Figure 4, and fits the data quite well: χ2=30.22, df=29, p=0.35, RMSEA=0.036 [0.000-0.107], CFI=0.995. The good fit of the full model suggests that the observed covariance pattern in our data is consistent with the statistical constraints imposed by the model, and allows us to further investigate the relations between the cognitive factors and the neural variables. The full model for Item Memory (Fig. 5), also fits the data well: χ2=21.09, df=29, p=0.5, RMSEA=0.008 [0.000-0.098], CFI=0.999. While both models share a number of structural and functional inputs, there are a number of unique inputs to each model (discussed below).

**Figure 4.**
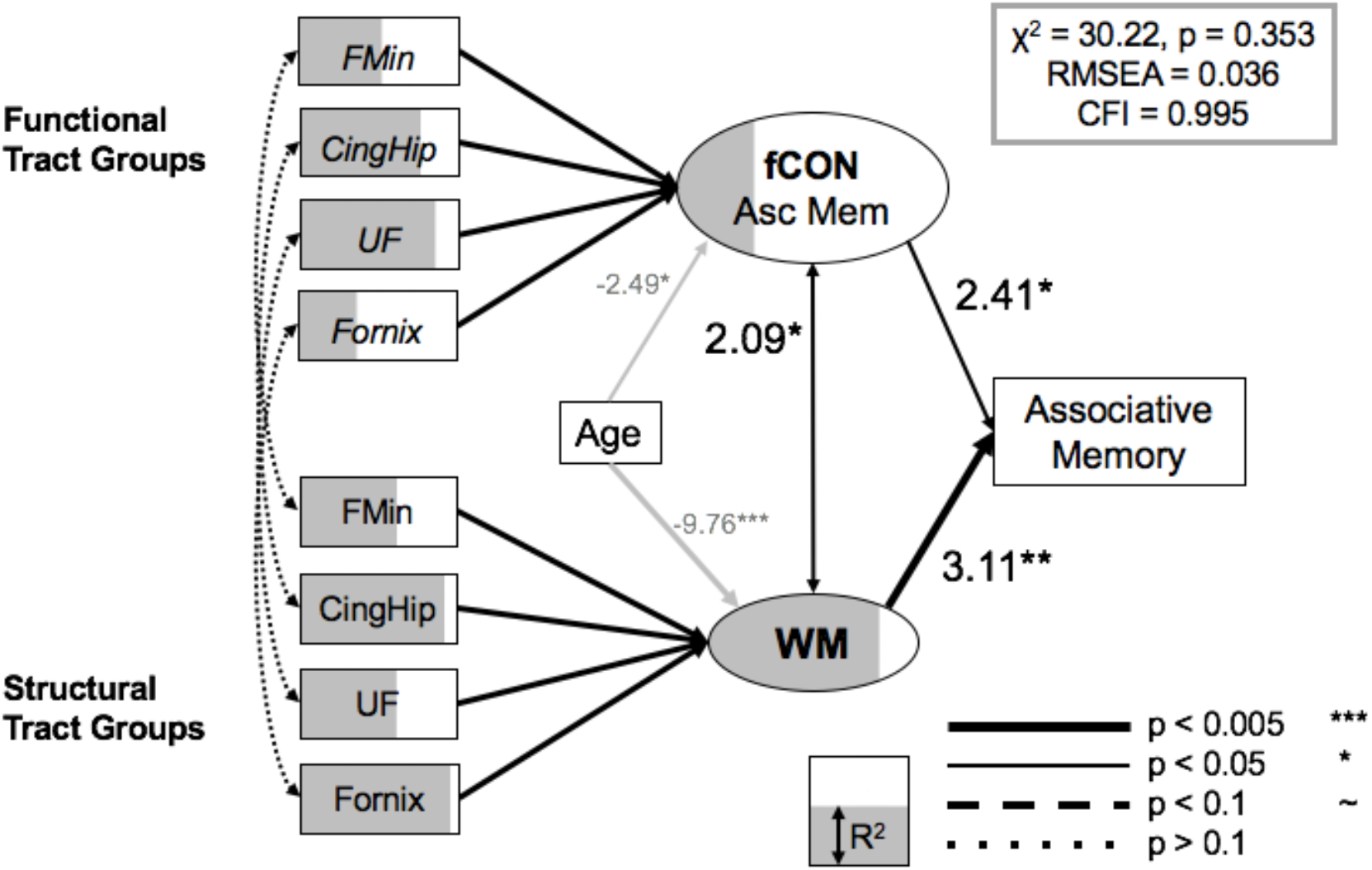
Full model, Associative Memory. Significant paths outlined and R^2^ is represented as the degree of shading of the variables. Brain measures only have paths to a corresponding CTG in the other modality, or to the appropriate LV. Notably, no tract-specific linkages between functional and structural information (left side of SEM). Note: CINGhip – ventral leg of the cingulum; FMin – forceps minor; UF – uncinate fasciculus.

**Figure 5.**
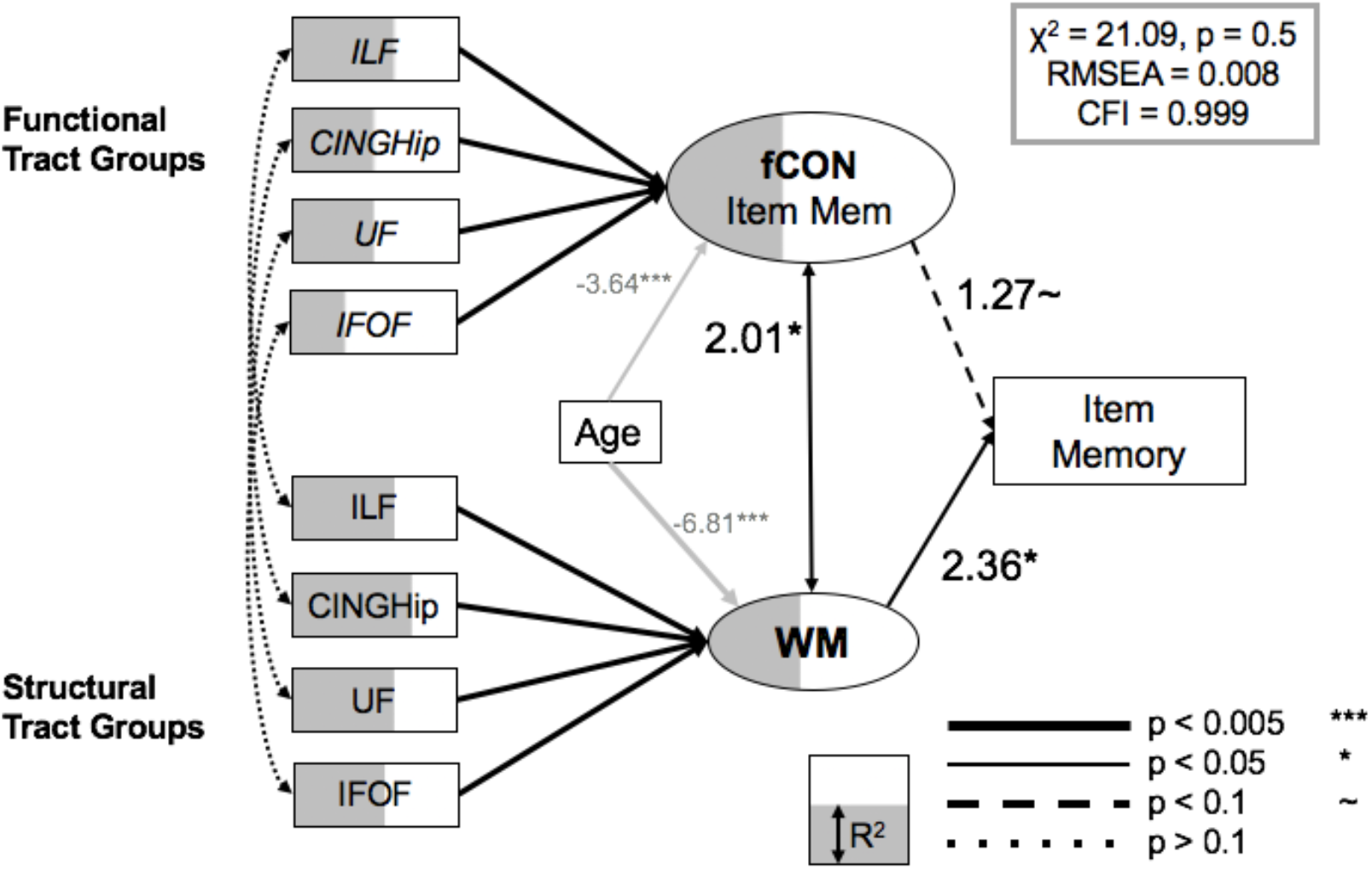
Full model, Item Memory. Significant paths outlined and R^2^ is represented as the degree of shading of the variables. Brain measures only have paths to a corresponding CTG in the other modality, or to the appropriate LV. Notably, just as in the Associative SEM above, no tract-specific linkages between functional and structural information. Note: CINGhip – ventral leg of the cingulum; IFOF – inferior-fronto-occipital fasciculus; ILF – inferior longitudinal fasciculus; UF – uncinate fasciculus.

#### Tract-specific contributions

We relied on the LASSO regularization to simplify our model and improve model fit; this technique also implicitly provides a means of identifying the specific tract groups that contribute to memory performance. As noted in the Methods, we constrained model terms to include both structural and functional information pairs for each CTGs, such that we could continue to make explicit hypotheses about structural-functional relationships in our final models. In the Associative memory model, the uncinate fasciculus, fornix, forceps minor, and hippocampal segment of the cingulum each contributed to the overall model (all R^2^ > 0.61/0.35 for structural/functional information, resp.). Furthermore, the inclusion of structural and functional information from the forceps minor of the corpus callosum, is in line with previous findings that suggest an important role for prefrontal WM in episodic memory functioning in older adult populations (Davis et al., 2009; Kennedy and Raz, 2009).

In contrast, the final Item Memory model relied on paired structural and functional information from the inferior longitudinal fasciculus, the hippocampal segment of the cingulum, the uncinate fasciculus, and the inferior fronto-occipital fasciculus (all R^2^ > 0.31/0.34 for structural/functional information, resp.). This result is consistent with the qualitative interpretation that Associative memory relies on fronto-temporal regions, while item memory shows a greater dependence on systems base solely within the temporal lobe (Glisky et al., 1995; Spaniol and Grady, 2012). Lastly, while the latent variable capturing the variance in overall success-related functional connectivity did not demonstrate a significant path to behavioral performance on the Item memory task (z = 1.27, p = 0.11), these functional CTGs nonetheless contributed to the overall model fit; a separate model removing the link from the fCON LV to Item Memory showed a significant reduction in model fit (Δχ^2^=39.15, Δdf=1, p < 0.01).

**Table 2.**
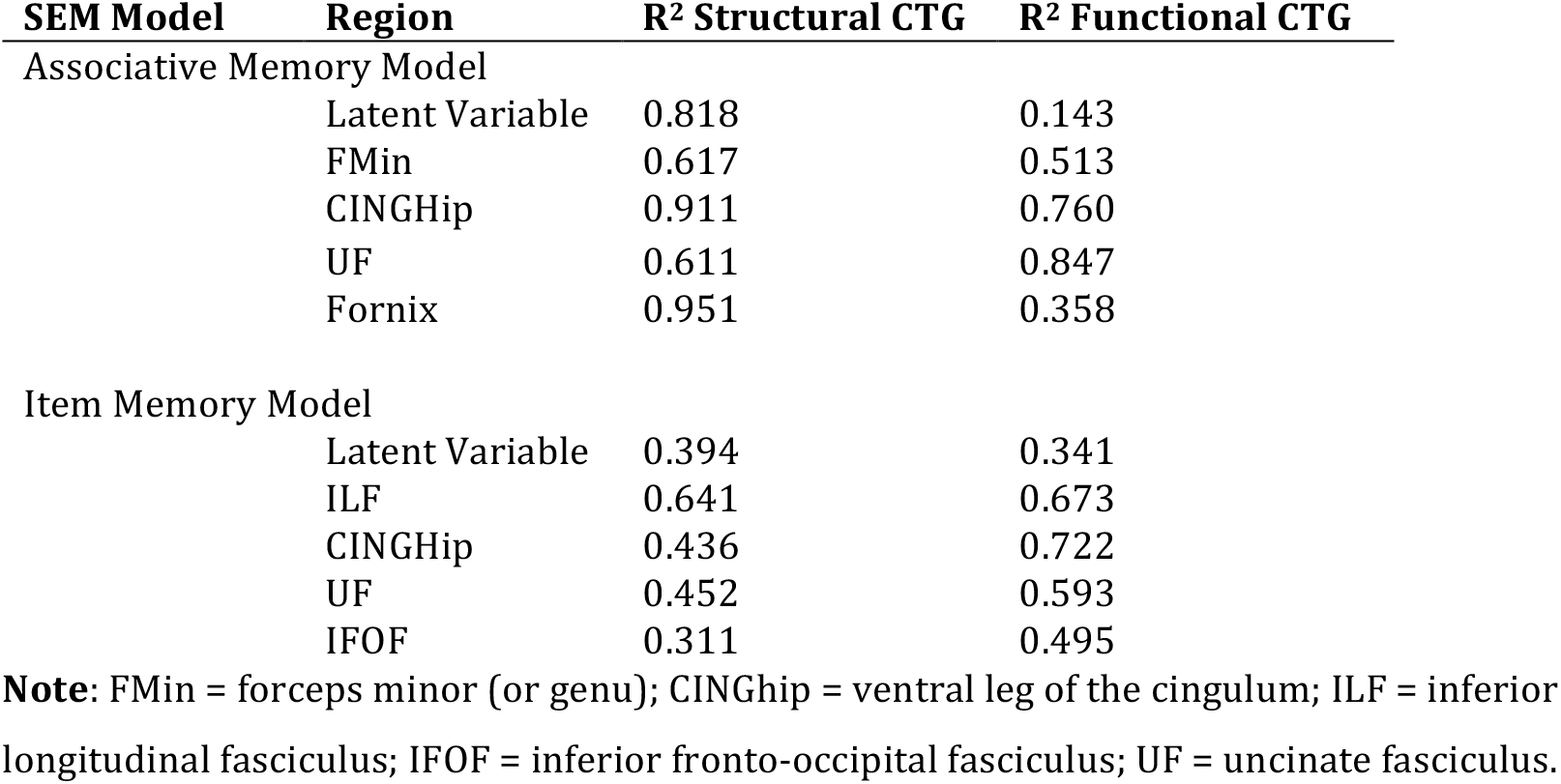
CTG-Specific Contributions.

#### Effects of Age

Our last question was whether the connectivity measures examined herein captured the effect of age on memory; in our full model above, Age effects on fCON and WM were highly significant in both Associative (Age➔fCON: z = −2.49; Age➔WM: z = −9.76) and Item Memory (Age➔fCON: z = −3.64; Age➔WM: z = −6.81) models. This result is unsurprising given the effects of age observed in Table 2. Nonetheless, the role of Age in explaining age-related changes in Associative and Item Memory is more adequately characterized within the full SEM. We therefore tested an alternative model in which Age effects were eliminated from the model. The overall fit was significantly worse in a model in which age were included, but paths from age to brain factors were fixed at zero, in both the Associative Memory (Δχ^2^= −89.21, Δdf=2, p < 0.001) and Item Memory models (Δχ^2^= −66.26, Δdf=2, p < 0.005). This result, unsurprisingly, demonstrates that chronological age captures a significant portion of variance in the model, and has a strong influence on the latent variables for structural and functional connectivity.

Finally, given the strong relationship between age and white matter measures, it is possible that structural or functional connectivity information in other tract groups capture additional age-related variance beyond the 4 CTGs that survived LASSO regularization for each model. We therefore performed a more unconstrained SEM with all 8 CTGs were included, and examined the change in model fit, as above. In the Associative Memory model, there was a drastic reduction in model fit, whether we included all 8 structural CTGs (Δχ^2^=64.31, Δdf=1, p < 0.005), functional CTGs (Δχ^2^=25.65, Δdf=1, p < 0.01), or both (Δχ^2^=99.79, Δdf=2, p < 0.001). Similarly, the ability of our model to capture age effects was not simply a function of any four CTGs. Model fits using the same number but different sets of CTGs (with equivalent Age effects) were similarly poor in model fit (all RMSEA > 0.3). Equivalent results were obtained in the Item Memory SEM. These results reinforce the importance of our regularization technique, and suggest that the additional information would not improve the inferences made in the above models. Given the strong collinearity with age in these additional terms (especially for all white matter regions), this null result suggests that unique role these regions in mediating the age-related declines in Associative and Item memory, and that these declines cannot be attributable to a general aging factor across all white matter tracts.

## Discussion

By combining multiple behavioural, demographic, and brain measures from a large sample of younger and older adults, we provide evidence that age-related differences in Associative and Item Memory are dissociable by their functional and structural connectivity profiles. In our best-fitting model, individual CTGs based on canonical fiber systems make independent contributions to both forms of memory. First, we found that the relationship between structural and functional connectivity information was best characterized by an intermediate level of relationship. Although no specific Tract Groups were linked in their corresponding fCON and WM values (e.g., fCON—WM linkage for the UF, ILF, etc.), a general WM—fCON relationship between latent variables inferred from these tract-specific measurements was significant in both Associative and Item memory SEMs. Second, we found that both WM and fCON make independent contributions to Associative Memory performance, while only WM influenced behaviour in the Item Memory model. Lastly, age-related influences on our model were much stronger for WM than fCON, but Age was an essential component of the full model. Our results therefore demonstrate that age-related declines in memory are unlikely to be driven by a single fiber system or a single data type, but emerge as a confluence of functional and structural changes in multiple anatomically connected systems.

### Structure—Function Relationships

Multiple evidences have shown that brain topology (i.e., structure) supports fluid dynamics (i.e., function), and that brain dynamics in turn reinforce structure via synaptic plasticity. In a very influential work Honey et al. (2007) showed that this relationship is highly dependent on the characteristics of the functional data used to test this relationship, including the timescale, local clustering, and brain state. Thus, understanding the precise relationship between these two forms of connectivity is still challenging, both methodologically (*how to compare these forms of data?*) and theoretically (*what mechanisms link these two forms of information?*). In the present analysis, we put forth a robust method to approach the former problem at a large-scale level of brain organization by addressing their mutual relationships during memory functioning, through a common connectivity key. By summarizing the structural (FA based on tractography streamlines) and functional (Spearman’s rho based on task-related PPI) relationships between pairs of regions which are connected by a canonical tract group (e.g., the uncinate fasciculus), our use of CTGs integrates structural and functional connectivity information within a common anatomical framework, by constraining functional connections to known anatomy. Furthermore, this strategy helps to link empirical results obtained via adjacency matrices—a now common basis for most graph-theoretical approaches to characterizing aging brain networks—with clinically-minded approaches centered on canonical fiber systems.

In our best fitting model for Associative or Item memory, no tract-specific linkages between structural and functional connectivity information for the same CTG reached significance, while latent variables for structural and functional connectivity did show a significant association. When setting this LV-LV pathway to zero, model fit significantly decreased (Δχ^2^= 9.78, Δdf=1, p < 0.016). It is worthwhile to note that, outside the SEM framework, functional and structural CTGs were reliably correlated across subjects (Table 1; all but one CTG r > 0.21, even after adjusting for age). Taken together, these results suggest that the relationship between structural and functional connectivity estimates may be best characterized on an intermediate level. Many of the age-related changes to white matter may appear to manifest as global changes across different major white matter tracts (Penke et al., 2010), and driven by causal factors that affect white matter, such as small vessel disease, myelin depletion, or iron accumulation. Nonetheless, a growing, model-based literature is emerging that suggests that a more constrained set of critical white matter fiber systems (forceps minor, cingulum, uncinate fasciculus) provides the best fit for models seeking to explain age-related changes in and attention, memory and processing speed (Voineskos et al., 2010; Lovden et al., 2013; Kievit et al., 2016).

### Tract-specific Effects on Associative and Item Memory

The value of such debates rests on the reliability of the anatomical specificity used in creating structural and functional connectivity values to predict age-related changes in behavior. Our result suggests an *intermediate* conclusion to the general vs. specific debate: while specific structural-functional linkages for a specific fiber tract do not drive the success of a model of Associative or Item memory, there are nonetheless a subset of specific tract groups which provide the best fit to these data. We found that the structural and functional connectivity based on the fornix had a selective positive influence for Associative, but not Item memory, consistent with theoretical and empirical results supporting the role of this structure in associative retrieval (Aggleton and Brown, 1999; Antonenko et al., 2016). The fornix is a key white matter tract of the medial temporal lobe memory system, interconnecting the hippocampal formation with subcortical structures in the basal forebrain and diencephalon. There is evidence of altered WM microstructure in the fornix in healthy older adults (Persson et al., 2006; Antonenko et al., 2016), and measures of fornix microstructure may be useful in detecting early/preclinical AD stages (Nowrangi and Rosenberg, 2015). The finding that the uncinate fasciculus is implicated in both our Associative and Item memory SEMs is consistent with evidence linking this tract to age-related decline in memory functioning across a wide array of tasks, including visual object location (Metzler-Baddeley et al., 2011), color-picture associations (Lockhart et al., 2012), working memory (Burzynska et al., 2013) and verbal learning (Lancaster et al., 2016). Similarly, the Item memory-specific role of the IFOF fits well with electrostimulation-based studies which have shown semantic paraphasias in response to (disruptive) stimulation of this fiber system (Duffau et al., 2005).

Interestingly, we found that the genu, or forceps minor of the corpus callosum (FMin in our models), which connects left and right prefrontal cortex, contributed significantly to Associative, but not Item Memory, a finding which is consistent with the assumption that Associative memory is more dependent on PFC-mediated functions (Shimamura, 1995), and that bilateral PFC activity may serve a compensatory role in age-related decline (Cabeza, 2002). We and others have found that the integrity of the genu predicts behavior on a range of episodic memory (Davis et al., 2009; Henson et al., 2016) and executive function tasks (Kievit et al., 2014; Kievit et al., 2016) in elderly populations. The dissociation in associative versus item memory performance (Glisky et al., 1995), and is consistent with abundant evidence that the PFC is more critical for associative than item memory performance in aging populations (Duarte et al., 2006; Old and Naveh-Benjamin, 2008; Spaniol and Grady, 2012; Leshikar and Duarte, 2014), and that reliance on structural and functional connectivity in the PFC may be a means of counteracting observed associative deficits (Naveh-Benjamin et al., 2003; Dennis et al., 2008).

### Effects of Age

Age effects on structural connectivity revealed with DWI are regionally diverse and typically show an anterior-to-posterior gradient of age-related decline (Sullivan et al., 2006; Davis et al., 2009). Consistent with this evidence, we observed strong declines in FA across nearly all structural connectivity groups, or CTGs (Table 2). While a number of studies have found single correlation (Kennedy and Raz, 2009) or mediation patterns (Madden et al., 2010; Oberlin et al., 2016) that help to explain how these white matter structures mediated cognitive decline, our analysis advances on these approaches by considering all of these regions simultaneously, within a statistically rigorous framework. We found that successful Associative memory was associated with structural and functional connectivity in a set of connecting within or between frontal regions: the forceps minor, UF, CINGhipp, and the fornix (Fig. 4), even when all brain measures were adjusted for chronological age. In contrast, an overlapping, but more ventral set of regions, including the inferior longitudinal fasciculus, helped to predict successful Item memory performance (Fig. 5).

With respect to functional connectivity, that connectivity groups connecting either within (forceps minor) or with (UF, IFOF) the PFC is consistent with the general observation that older adults show higher levels of PFC activity across all a range of cognitive tasks (Grady, 2012). Our findings of age-related increases in task-related frontotemporal functional connectivity during successful Associative memory provide strong evidence for a compensatory mechanism by demonstrating a reliance on PFC connectivity from 3 major inferior temporal white matter fibers: the FMin, UF, and fornix. Our connectivity measure was based upon task-related data, and furthermore used an explicit contrast between Hits > Misses, in order to isolate connectivity related to successful memory functioning. Our result is also consistent with a handful of studies which have identified an increased reliance on frontotemporal connectivity to maintain associative memory function (Dennis et al., 2008; Spaniol and Grady, 2012), and this finding more generally supports the idea that frontotemporal interactions support associative or source memory performance (Backus et al., 2016). Despite these steps forward, it is important to remember that our data are cross-sectional. The nature of these data therefore makes it difficult to distinguish true effects of “age” from cohort effects related to year of birth. Indeed, longitudinal studies (Ronnlund et al., 2005; Nyberg et al., 2010) have shown important differences between the effects of age and effects of birth year. In order to establish a more robust model of the relationship between structural and functional dynamics, it would be necessary to follow people over time, and establish what causal factors (critical developmental periods, nutrition, cardiovascular fitness, etc.) influence this relationship.

### Using SEM to investigate network connectivity

The differential sensitivity of associative and item memory to both structural and functional connectivity supports models in which these types of brain measures constitute dissociable indices of brain health. Moreover, our results argue for the importance of general factors predicting age-related declines in brain health, over tract-specific linkages to memory functioning. These are some of the issues addressed by turning to SEM as a means of fitting the pattern of covariance between our connectivity measures (rather than their mean values), and using this information to predict the outcomes for behavioral performance (Voineskos et al., 2010). Moreover, we can use model selection to weight parsimony versus explanatory power of competing connectivity models, testing their relative degree of support in our sample. Here we sought to develop a single, robust method for combining structural and functional connectivity information within an anatomically informed framework. Furthermore, we used a LASSO penalty to estimate to accurately and efficiently estimate a model that is parsimonious. Thus, our theoretically-driven approach uses multiple robust statistical approaches to reduce the complexity of connectivity information inherent in multidimensional connectomes, and helps to resolve a sensible model of Associative and Item memory using multiple anatomically-based predictors.

### Conclusions

To summarize, based on the evidence that structural and functional networks from our CTG analysis, we have shown that age-related changes in associative—but not item memory—are critically dependent on the linkages between structural and functional connectivity tract groups. Usually, an implicit assumption is that the structure of the network is observable, and inference of the underlying structure of the connected system can be based on diffusion tractography techniques. Our results test this assumption explicitly, by using an analytical method that puts structural and functional connectivity information on equal footing. These results provide further insights into the interplay between structural and functional connectivity patterns, and help to elucidate their relative contribution to age-related changes in associative memory performance.

